# Alternative splicing controls pan-neuronal homeobox gene expression

**DOI:** 10.1101/2024.07.16.603806

**Authors:** Eduardo Leyva-Díaz, Michael Cesar, Karinna Pe, Jose Ignacio Jordá-Llorens, Jessica Valdivia, Oliver Hobert

**Affiliations:** Department of Biological Sciences, Columbia University, Howard Hughes Medical Institute, New York, NY, USA; Department of Developmental Neurobiology, Instituto de Neurociencias (CSIC-UMH), Alicante, Spain

## Abstract

The pan-neuronally expressed and phylogenetically conserved CUT homeobox gene *ceh-44/CUX* orchestrates pan-neuronal gene expression throughout the nervous system of *C. elegans.* As in many other species, including humans, *ceh-44/CUX* is encoded by a complex locus that also codes for a Golgi-localized protein. How gene expression from this complex locus is controlled and, in *C. elegans*, directed to all cells of the nervous system has not been investigated. We show here that pan-neuronal expression of CEH-44/CUX is controlled by a pan-neuronal RNA splicing factor, UNC-75/CELF, the *C. elegans* homolog of vertebrate CELF proteins. UNC-75/CELF temporally and spatially specifies the production of an alternative, CEH-44/CUX homeobox gene-encoding transcript from a ubiquitously expressed gene locus, which also produces a Golgi apparatus-localized golgin protein, CONE-1 (“**<u>C</u>**ASP **<u>o</u>**f **<u>ne</u>**matodes”). During embryogenesis the *cone-1/ceh-44* locus exclusively produces the Golgi-localized CONE-1/CASP protein in all tissues, but upon the onset of postmitotic terminal differentiation of neurons, UNC-75/CELF induces the production of the alternative CEH-44/CUX CUT homeobox gene-encoding transcript, exclusively in the nervous system. Hence, UNC-75/CELF-mediated alternative splicing not only directs pan-neuronal gene expression, but also excludes a phylogenetically deeply conserved golgin from the nervous system, paralleling surprising spatial specificities of another golgin that we describe here as well. In summary, our findings provide novel insights into how all cells in a nervous system acquire pan-neuronal identity features.

## INTRODUCTION

The underlying basis for the uniqueness of any animal’s nervous system is the enormous diversity of cell types that populate this organ. Neuronal gene expression programs can be broken down into two separate, parallel acting routines: Genes that are selectively expressed in specific types of neurons are coordinately controlled by combinations of master regulatory transcription factors, called terminal selectors (Hobert 2016). Loss of terminal selectors therefore results in the loss of the cell type-specific identity of a neuron class. In contrast, loss of terminal selectors does not affect the expression of pan-neuronal genes (e.g. neuropeptide processing machinery, synaptic vesicle proteins, etc.), demonstrating the existence of parallel gene regulatory programs that define the pan-neuronal features of a neuron (Hobert et al. 2010).

We have recently uncovered the identity of pan-neuronal gene expression regulators. We found that an entire subfamily of homeobox genes, the CUT homeobox genes, operates in a redundant and dosage-dependent manner to control pan-neuronal gene expression (Leyva-Diaz and Hobert 2022). Several of the CUT homeobox genes are ubiquitously expressed, but two of them, *ceh-44/CUX* and *ceh-48/ONECUT*, are restricted to the nervous system, apparently increasing CUT gene dosage above a level sufficient to control pan-neuronal target gene expression. However, how *ceh-44/CUX* and *ceh-48/ONECUT* expression is restricted to the nervous system, is not known. In this paper, we describe an unanticipated regulatory strategy that drives expression of *ceh-44/CUX* in the nervous system. This regulatory strategy also sheds light on a highly unusual and mysterious gene locus that is present throughout the animal kingdom, from worms to humans (Burglin and Cassata 2002; Takatori and Saiga 2008).

More than 20 years ago, in a search for proteins involved in trafficking at the Golgi apparatus, Gillingham and colleagues noted that a phylogenetically deeply conserved, Golgi-localized protein is encoded by a genetic locus that is also capable of producing a homeodomain protein (Gillingham et al. 2002)(**Fig.1A**). The Golgi localized isoform of this locus, called CASP, is conserved from yeast to mammals and functions as a so-called “golgin”, a group of Golgi-membrane proteins unrelated by primary sequence that are involved in vesicle transport along the Golgi apparatus (**Fig.2A**)(Munro 2011; Lowe 2019). The homeobox containing isoform of this locus also encodes a deeply conserved protein, the CUT homeodomain protein CUX1, the vertebrate ortholog of *C. elegans* CEH-44 protein. The structural organization of this locus is such that CASP and CUX1 share the N-terminal region, while their specific domains differ downstream (**Fig.1A**).

**Figure 1:**
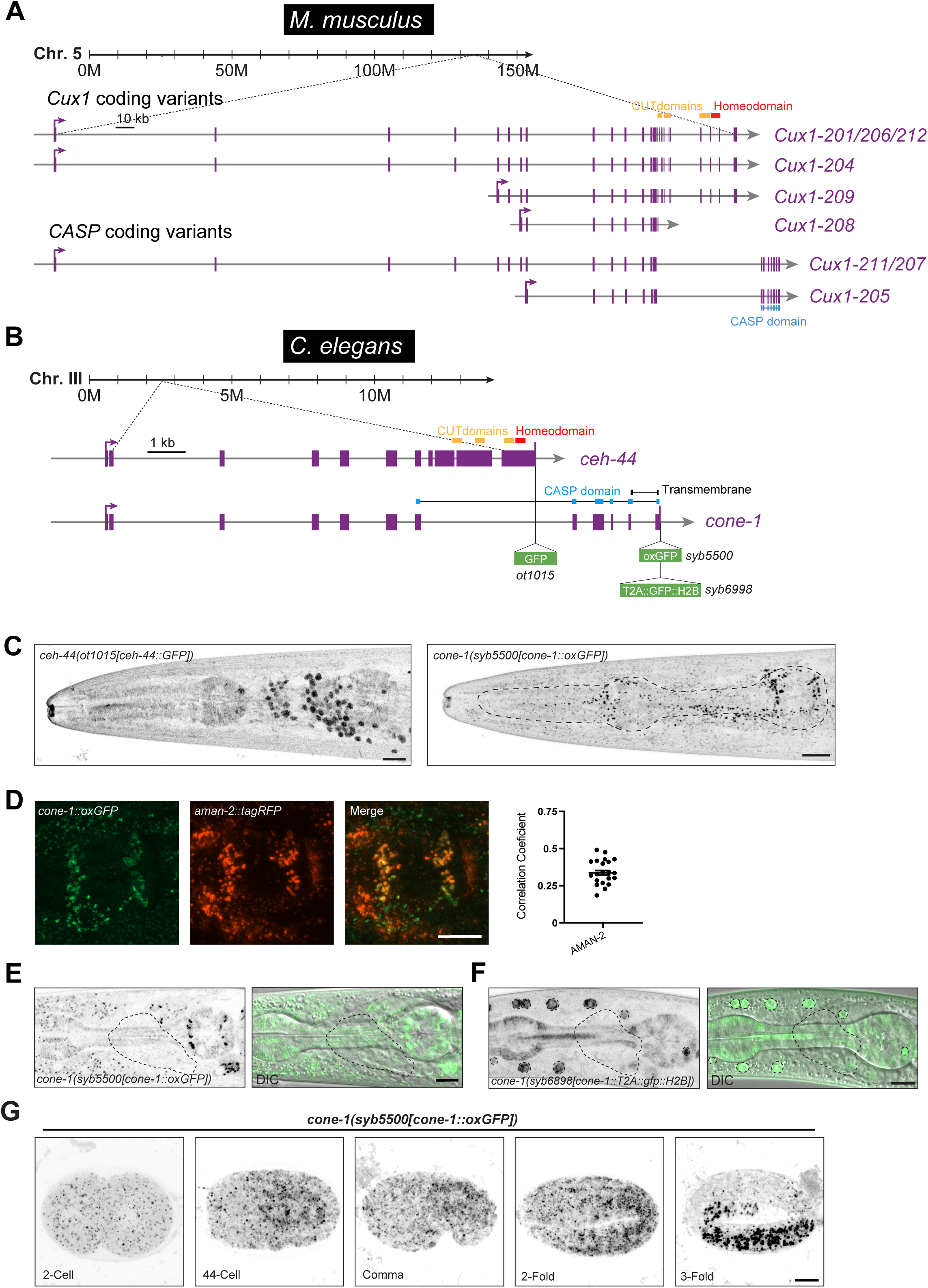
*cone-1/ceh-44* locus structure and expression pattern. (**A**) Schematic illustration of the mouse *Cux1* locus showing the predicted transcripts coding for Cux1 and CASP (Ensembl release 112 (Harrison et al. 2024)). (**B**) Schematic representation of the *C. elegans cone-1/ceh-44* gene locus showing the location of the GFP insertions, homeodomain, CUT and CASP domains. (**C**) Reporter expression in L4 animals (head, lateral views), left *ceh-44(ot1015[ceh-44::gfp])* showing pan-neuronal nuclear expression, and right *cone-1(syb5500[cone-1::oxGFP])* showing broad puncta. (**D**) Reporter expression overlap in L4 animals (head, lateral view, posterior bulb pharynx) of *cone-1(syb5500[cone-1::oxGFP])* and *aman-2prom::tagRFP[pwIs1022]*. Pearson’s correlation coefficient measurements for colocalization of CONE-1::GFP with AMAN-2::tagRFP (right). (**E-F**) Expression of *cone-1(syb5500[cone-1::oxGFP])*(**E**) and *cone-1(syb6898[cone-1::T2A::gfp::H2B])* (**F**). Green/DIC channels merge showing expression excluded from the nervous system (right, single Z-slice). Lateral views of the worm head at the L4 stage are shown. Dashed contour outlines the nerve ring region. (**G**) Temporal expression analysis in *cone-1(syb5500[cone-1::oxGFP])* across different embryonic stages: 2-Cell, 44-Cell, Bean, 2-Fold, 3-Fold. Scale bars 10 μm.

**Figure 2:**
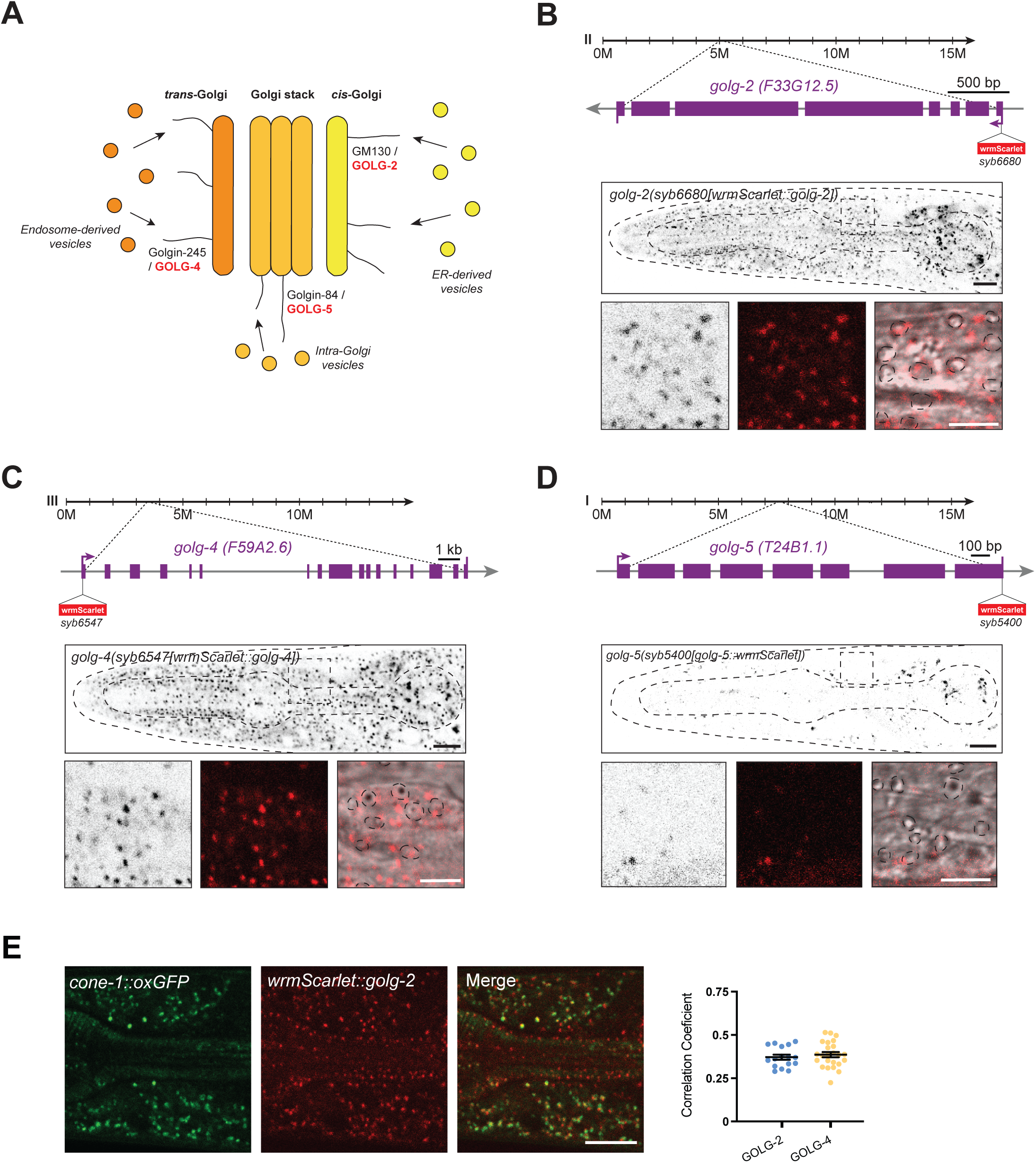
*C. elegans* golgin proteins show ubiquitous and cell-type specific expression patterns. (**A**) Different golgins localize to different Golgi compartments. *C. elegans* orthologs that are tagged in this study are labeled in red. Schematic adapted from (Lowe 2019). (**B-D**) Schematic representations of the golgin loci showing wrmScarlet insertion locations. Lateral head images of L4 animals showing expression of *golg-2(syb6680[wrmScarlet::golg-2])* (**B**), *golg-4(syb6547[wrmScarlet::golg-4])* (**C**), and *golg-5(syb5400[golg-5::wrmScarlet])* (**D**) (single Z-slices). Magnified images of area between bulbs shown to the right, shown as black and white, red, and merge with DIC channel. (**E**) Reporter expression overlap in L4 animals of *cone-1(syb5500[cone-1::oxGFP])* and *golg-2(syb6680[wrmScarlet::golg-2])* (head, lateral view). Pearson’s correlation coefficient measurements for colocalization of CONE-1::GFP with golgin proteins (right). ER, endoplasmic reticulum. Scale bars 10 μm (5 μm in zoomed images in **B-D**).

A substantial number of questions about this locus have remained unsolved: Are those two isoforms really the product of alternative splicing? Are the alternative transcripts generated in an overlapping or mutually exclusive manner? How is this alternative splicing regulated? And why are these two gene products coupled in this manner? Here, we provide answer to several of these question using the conserved, golgin/homeobox gene locus structure in *C. elegans* as a model (**Fig.1B**). We place the regulation of this locus into the context of control of pan-neuronal gene expression, finding that in *C. elegans* the *ceh-44/CUX* isoform of this locus is produced exclusively in the nervous system, via the action of a pan-neuronal expressed, alternative splicing factor, UNC-75, the *C. elegans* homolog of the vertebrate CELF splicing regulators. This splicing pattern results in the exclusion of the alternative, *cone-1/CASP-*encoding transcript, that codes for the *C. elegans* homolog of the CASP golgin, from the nervous system. We find that another *C. elegans* golgin protein also displays cell type specification expression patterns, thereby uncovering cellular diversity of composition and, possibly, function of the Golgi apparatus.

## RESULTS

### The *cone-1/ceh-44* locus generates two differentially expressed and localized proteins: the Golgi-localized CONE-1/CASP and the nuclear CUT homeodomain CEH-44/CUX

CUT homeobox genes are highly conserved transcription factors with orthologs ranging from nematodes to mammals (Burglin and Cassata 2002). In *C. elegans*, *ceh-44/CUX* represents the only member of the CUX subfamily of CUT homeobox genes, characterized by the presence of multiple CUT domains (Burglin and Cassata 2002). In vertebrates, the *Cux1* locus produces an alternative transcript that encodes CASP (<u>C</u>ux1 <u>A</u>lternatively <u>S</u>pliced <u>P</u>roduct), a single pass transmembrane protein that localizes to the Golgi apparatus (**Fig. 1A**)(Gillingham et al. 2002). Similarly, the *C. elegans* locus encoding *ceh-44/CUX* produces the ortholog of the vertebrate CASP, which we named “*cone-1*” (<u>C</u>ASP <u>o</u>f <u>ne</u>matodes).

We have previously reported the pan-neuronal expression pattern of *ceh-44/CUX* by generating an endogenous GFP reporter via CRISPR/Cas9-mediated genome engineering (Leyva-Diaz and Hobert 2022). However, the expression and potential subcellular localization of *cone-1/CASP* has not been examined. To address this question, we CRISPR/Cas9-engineered a translational reporter allele, *cone-1(syb5500)*, by inserting GFP at the C-terminus end of CONE-1/CASP (**Fig. 1B**). Since based on its sequence, we predict CONE-1/CASP to be a transmembrane protein with its C-terminus portion present in the acidic environment of the Golgi apparatus, we employed a worm-codon optimized GFP form (oxGFP) engineered to maintain its structure under acidic conditions. We found that CONE-1::oxGFP is broadly expressed throughout the nematode body, displaying a marked punctate pattern, contrasting the nuclear pan-neuronal expression of the CEH-44/CUX isoform of this locus (**Fig. 1C**). Using an RFP marked Golgi protein, AMAN-2, we found an overlap of CONE-1/CASP and AMAN-2 localization, indicating that like vertebrate CASP, the nematode CONE-1 protein is a Golgi-resident protein (**Fig. 1D**).

CONE-1/CASP puncta are conspicuously absent from neuronal cells (**Fig. 1E**), hence displaying a mutually exclusive expression with *ceh-44/CUX*. To validate this observation, we generated a *cone-1/CASP* reporter allele, *cone-1(syb6998)*, in which we inserted a nuclear localized GFP reporter cassette at the 3’ end of the *cone-1/CASP* transcript, separated from CONE-1/CASP protein by an “ribosomal skip” T2A peptide (Ahier and Jarriault 2014). This nuclear-directed reporter allows for independent identification of sites of expression and validates exclusion of CONE-1/CASP protein expression from the nervous system (**Fig. 1F**).

Intriguingly, we found a notable difference in expression onset during embryonic development between CEH-44/CUX and CONE-1/CASP proteins. As we have previously described, CEH-44::GFP expression initiates promptly after neuronal birth at the embryonic comma stage (Leyva-Diaz and Hobert 2022). In contrast, we detect CONE-1/CASP expression, as assessed with the translational CONE-1::oxGFP reporter as early as the 2-cell stage, displaying a broad, seemingly ubiquitous expression pattern across all proliferative embryonic stages (**Fig. 1G**), followed by exclusion of CONE-1 and onset of CEH-44 expression in the maturing, postmitotic nervous system.

### Cell-type specificity of golgin expression

The neuron-excluded expression of CONE-1/CASP protein prompted us to test whether other golgin proteins may also display cell-type specific expression profiles. To this end, we considered *C. elegans* orthologs of three different golgin proteins described in other systems: GM130, golgin-245 and golgin-84, each labeling distinct Golgi compartments (GM130 *cis*-Golgi, golgin-84 Intra Golgi vesicles, and golgin-245 *trans*-Golgi) (Munro 2011; Lowe 2019) (**Fig. 2A**). We termed these orthologs *golg-2* (GM130 ortholog), *golg-4* (golgin-245) and *golg-5* (golgin-84). wrmScarlet, a worm codon-optimized version of mScarlet (El Mouridi et al. 2017) was inserted either right after the initiation codon of the gene (*golg-2* and *golg-4*) or right before the termination codon of the gene (*golg-5*) using CRISPR/Cas9 genome engineering (**Fig. 2B-D**). Punctate expression patterns were observed in each of the three strains. In non-neuronal cells, these punctae overlapped with CONE-1/CASP localization, corroborating the Golgi localization of these proteins (**Fig. 2E**). In all three cases, expression persists throughout all developmental stages into adulthood. Intriguingly, while GOLG-2 and GOLG-4 are broadly, if not ubiquitously expressed in the adult animal, GOLG-5 protein appears to be expressed only in a subset of cell types throughout the animal (**Fig. 2D)**. Therefore, we have uncovered here the first two examples of cell-type specific golgin protein expression, GOLG-5 and CONE-1.

To assess CONE-1/CASP protein function, we generated a *cone-1/CASP*-specific mutant allele through the insertion of a premature stop codon in the first *cone-1/CASP-*specific exon (**Fig. S1A**). No obvious developmental or behavioral defects were observed in this *cone-1/CASP* mutant allele. Moreover, we found that neither the expression of *ceh-44/CUX* nor its function, assessed by the expression of *ric-19*, a pan-neuronal gene controlled by *ceh-44/CUX* (Leyva-Diaz and Hobert 2022), are altered in animals lacking *cone-1/CASP* (**Fig. S1B-C**). The absence of *cone-1/CASP* phenotype aligns with the lack of phenotypes observed after knockout of the yeast homolog of CONE-1/CASP, called COY, or other golgins in mice or flies (Matsukuma et al. 1999; McGee et al. 2017; Park et al. 2022). Similarly, deletion or nonsense alleles of *golg-2*, *golg-4* and *golg-5* have been generated by the *C. elegans* knock-out consortia and appear viable, consistent with viable phenotypes of fly and mouse homologs of these genes.

### Deconstructing CONE-1 and CEH-44 coding regions

While *ceh-44/CUX* and *cone-1/CASP* probably serve different functions, they are predicted to share a large portion of their N-terminal region. However, the key CEH-44/CUX functional domains, the CUT and homeodomain DNA binding domains, are exclusively encoded within the *ceh-44/CUX* specific exons. Similarly, vertebrate *Cux1* and *CASP* also share many exons, and the CUT and homeodomain sequence are also encoded by *Cux1*-specific exons (**Fig. 1A**). To experimentally confirm predictions of gene structures and their cell-type specific expression, we first examined transcripts from the *cone-1/ceh-44* locus in available neuronal RNA-seq data (Leyva-Diaz and Hobert 2022). Confirming our reporter-based expression data, *cone-1/CASP* specific exons are absent in neuronal transcripts of this locus, *ceh-44/CUX* specific exons are well covered by neuronal RNA-seq reads (**Fig. 3A**). Intriguingly, coverage within the shared exons starts only in exon 4, indicating a potential alternative *ceh-44/CUX* start site (**Fig. 3A**). To test this notion, we introduced an early stop codon in exon 3 and found CEH-44::GFP protein localization and expression levels to be unaffected, while CONE-1::GFP protein expression was completely eliminated (**Fig. 3A-C**). Consistent with the production of a shorter CEH-44/CUX protein lacking exons 1-3, insertion of GFP via CRISPR/Cas9 into the first exon, right after the start codon (**Fig. 3A**; *syb5437* allele) produced Golgi-localized CONE-1::GFP expression, but no neuronal, nuclear CEH-44::GFP expression, further corroborating the use of an alternative start site for *ceh-44/CUX* (**Fig. 3D**, left).

**Figure 3:**
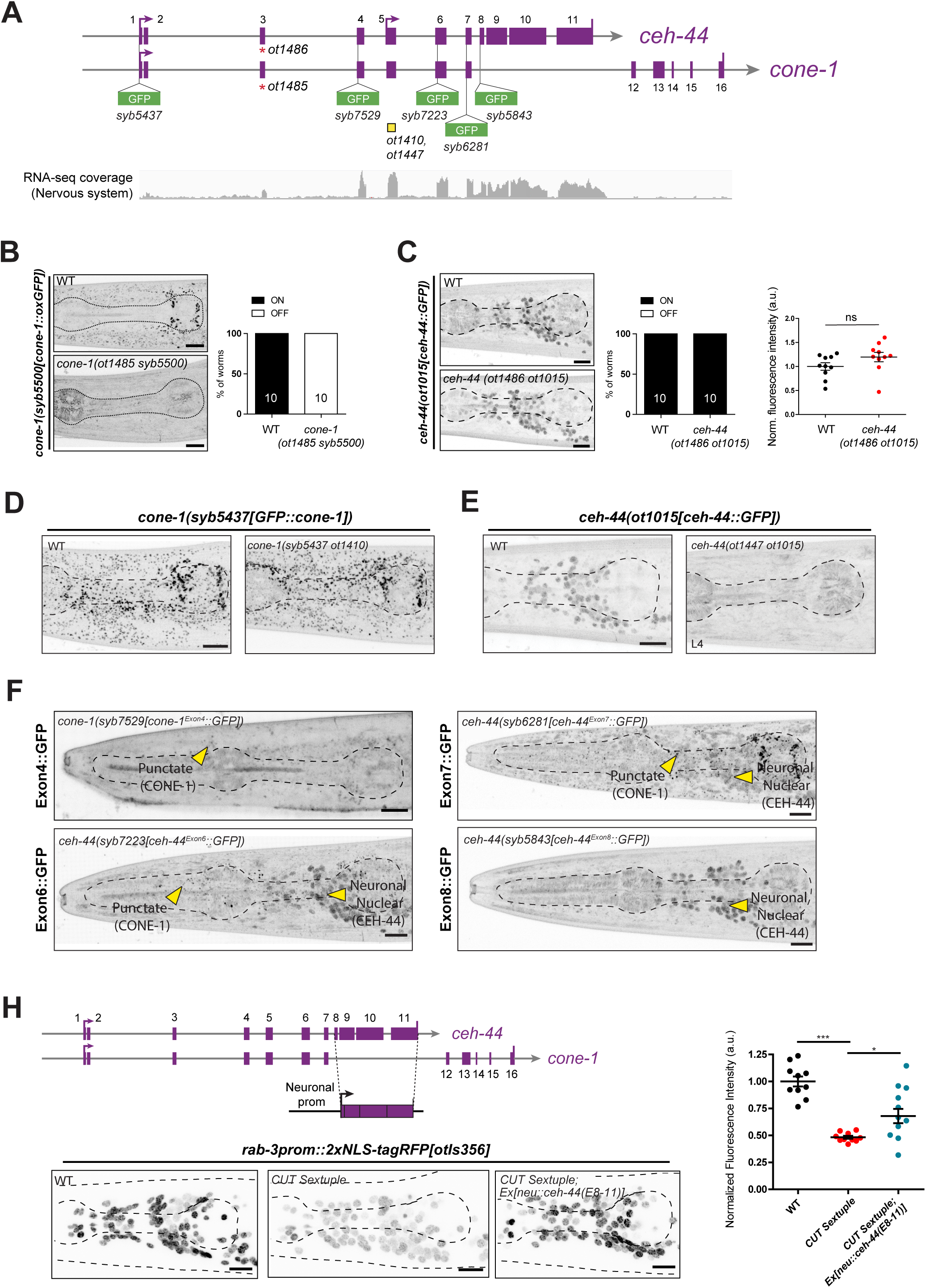
Revised locus structure: CEH-44/CUX and CONE-1/CASP present different start sites. (**A**) Schematic representation of *cone-1/ceh-44* gene locus showing GFP insertion locations and mutant alleles (yellow boxes indicate deletions, red asterisks indicate early stop codons). RNA-seq coverage track from isolated nervous system tissue shown at the bottom (RNA-seq, average, n = 3). (**B-C**) *cone-1(syb5500[cone-1::oxGFP])* (**B**) and *ceh-44(ot1015[ceh-44::gfp])* (**C**) reporter expression in L4 animals (head, lateral views) in wild-type (top) and mutants carrying an early stop codon in exon 3 (bottom). Note that the *ot1485* (**B**) and *ot1486* (**C**) mutant alleles are designed to maintain frame and harbor the same molecular lesion. Percentage of animals that express the reporter (ON/OFF) are indicated. CRISPR allele fluorescence intensity in head neurons was quantified in (**C**). The data are presented as individual values with each dot representing the expression level of one worm with the mean ± SEM indicated. Unpaired *t*-test, n = 10 for all genotypes. (**D-E**) *cone-1(syb5437[gfp::cone-1])* (**D**) and *ceh-44(ot1015[ceh-44::gfp])* (**E**) reporter expression in L4 animals (head, lateral views) in wild-type (left) and mutants carrying an exon 5 deletion (right). Note that the *ot1410* (**D**) and *ot1447* (**E**) mutant alleles are designed to maintain frame and harbor the same molecular lesion. See **Fig. S2C** for sequence details. (**F**) Reporter expression in L4 animals of *cone-1(syb7529[cone-1Exon4::gfp])* (top left)*, ceh-44(syb7223[ceh-44Exon6::gfp])* (bottom left)*, ceh-44(syb6281[ceh-44Exon7::gfp])* (top right)*, and ceh-44(syb5843[ceh-44Exon8::gfp])* (bottom right) (head, lateral views). Yellow arrows point to CONE-1/CASP puncta in *cone-1(syb7529[cone-1Exon4::gfp]),* both *cone-1/CASP* puncta and CEH-44/CUX neuronal nuclear expression. (**H**) Expression of *rab-3prom1::2xNLS-tagRFP[otIs356]* in wild-type (left), CUT sextuple mutant (middle), and CUT sextuple mutant rescue (pan-neuronal expression of *ceh-44/CUX* specific exons 8-11, *neu::ceh-44(E8-11)[otEx7645]*, “neu” = *ceh-48* promoter) (right). Images are lateral head views of L4 animals (max Z-projections). Quantification of fluorescence intensity in head neurons. Each dot represents the expression level within one worm with the mean ± SEM indicated. Wild-type data are represented with black dots, CUT sextuple mutant with red dots, and rescue with blue dots. One-way ANOVA followed by Tukey’s multiple comparisons test; *P < 0.05, ***P < 0.001. n ≥ 10 for all genotypes. ns, not significant; a.u., arbitrary units. Scale bars 10 μm.

To dissect the structure of CEH-44/CUX and CONE-1/CASP proteins further, we inserted *gfp* at different positions within the *cone-1/ceh-44* endogenous locus (**Fig. 3A, S2A**). First, CRISPR/Cas9-engineered *gfp* insertion in the *ceh-44/CUX* specific exon 8, the first exon of *ceh-44/CUX* specific exons, revealed pan-neuronal and nuclear CEH-44::GFP protein, as expected (**Fig. 3F**). *gfp* insertions within exons 6 or exon 7, which are shared between *cone-1/CASP* and *ceh-44/CUX,* resulted in simultaneous detection of CONE-1::GFP in the Golgi and CEH-44::GFP in neuronal nuclei (**Fig. 3F**). In contrast, only Golgi-localized CONE-1::GFP was observed when *gfp* was engineered into exon 4 (**Fig. 3F**).

Although exon 4 is covered in neuronal transcriptomic data (**Fig. 3A**), there is no potential start codon within exon 4, yet there are three start codons in exon 5, just prior to the GFP insertion within exon 6 (**Fig. S2B**). To test the importance of exon 5 and its embedded start codons, we engineered an in-frame deletion of most codons within this exon, including all potential start codons, using CRISPR/Cas9 (**Fig. 3A, S2C**). We found that in these animals CEH-44::GFP expression is completely abolished while CONE-1::GFP expression remained unaffected (**Fig. 3D-E**). Taken together, our data supports an updated *cone-1/ceh-44* locus structure with different start sites for each gene (**Fig. 3A**): *cone-1/CASP* initiation codon is present in exon 1, while the *ceh-44/CUX* initiation codon resides within exon 5. As we will describe in the next section, the initiation of the shorter *ceh-44/CUX* transcript may be an autoregulatory effect.

Finally, we sought to investigate the requirement of shared exons for *ceh-44/CUX* function. We have previously shown that the expression of pan-neuronal genes, such as the small GTPase *rab-3*, is significantly reduced in animals lacking all the neuronal CUT genes, including *ceh-44/CUX* (CUT sextuple mutants). Overexpression of a *ceh-44/CUX* construct containing exons 1 through 11 rescued pan-neuronal gene expression (Leyva-Diaz and Hobert 2022). Here, we found that a rescuing construct containing only *ceh-44/CUX*-specific exons (exons 8-11) also rescues *rab-3* expression (**Fig. 3H**), suggesting that *cone-1/ceh-44* shared exons might not be required for *ceh-44/CUX* function.

### Different regulatory elements control *ceh-44/CUX* pan-neuronal expression pattern

After defining the complex structure and expression of the *cone-1/ceh-44* locus, we sought to deconstruct its *cis-*regulatory architecture. To this end, we constructed reporter gene fusions that interrogate the *cis-*regulatory content within both upstream and intronic regions of the *cone-1/ceh-44* locus. First, we generated a reporter construct that contains 2kb upstream of the primary *cone-1/ceh-44* transcript. We found this reporter to drive expression broadly, both in neuronal and non-neuronal cells (**Fig. 4A**). To corroborate that this transcript is indeed produced in all cells, we engineered an endogenous reporter for this upstream region by inserting a *gfp::H2B::SL2* cassette just before the *cone-1/CASP* initiation codon (**Fig. S3A**). Due to the trans-splicing driven by the bicistronic SL2 linker (Spieth et al. 1993; Huang et al. 2001), this reporter served as a GFP transcriptional reporter for the *cone-1/ceh-44* upstream promoter. Expression analysis across different embryonic and larval stages unveiled widespread, ubiquitous expression in both neuronal and non-neuronal cells, commencing at the 2-cell embryo stage and persisting through adulthood (**Fig. S3B**). These observations suggest that this upstream region may function as an initiation and maintenance regulatory element for the locus.

**Figure 4:**
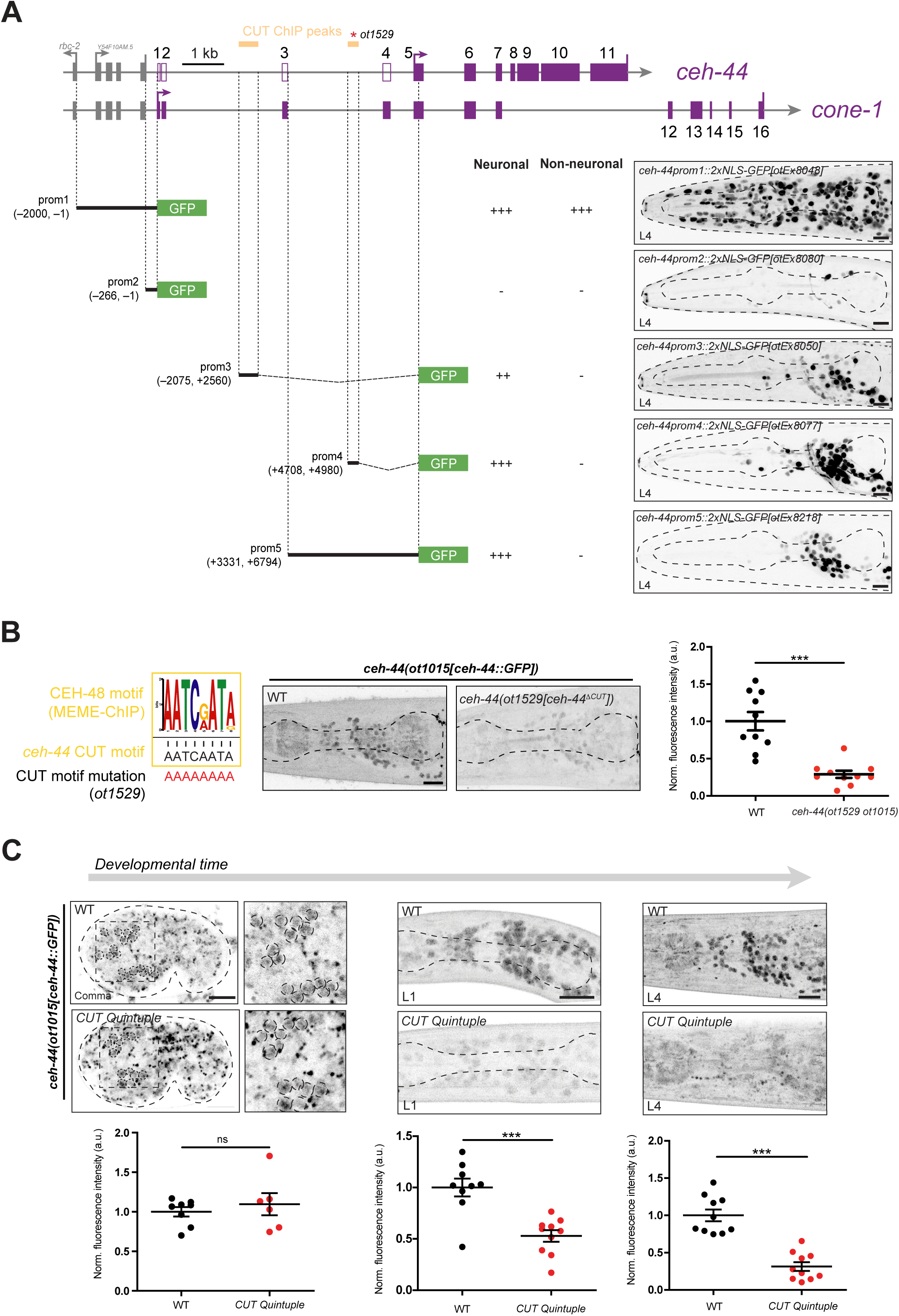
CUT homeobox genes maintain *ceh-44/CUX* expression. (**A**) Schematic representation of *cone-1*/*ceh-44* locus *cis*-regulatory analysis. CEH-48 and CEH-38 peaks found in the ChIP-seq datasets are displayed in yellow. Promoter 1-5 designs are displayed with coordinates, neuronal and non-neuronal expression is described as either (-) no expression, (++) broad expression, or (+++) pan-neuronal/ubiquitous expression. (**B**) *ceh-44(ot1015[ceh-44::gfp])* reporter expression in wild-type (left) and mutant animals carrying a mutation in the CUT homeodomain binding site within the intron 3 CUT ChIP peak (*ot1529* allele, right). The consensus binding motif for CEH-48 and CEH-38, extracted from the ChIP-seq datasets using MEME-ChIP (Machanick and Bailey 2011) are shown on the left. (**C**) *ceh-44(ot1015[ceh-44::gfp])* reporter expression at the comma embryonic stage (left), larval L1 stage (middle) and larval L4 stage (right) in wild-type (top) and CUT quintuple mutants (bottom). Quantification of CRISPR allele fluorescence intensity in head neurons (**B** and **C**). The data are presented as individual values with each dot representing the expression level of one worm with the mean ± SEM indicated. Wild-type data are represented with black dots and mutants with red dots. Unpaired *t*-test, ***P < 0.001. n ≥ 6 for all genotypes. Images show lateral views in all panels (L4 stage in **A** and **B**). ns, not significant; a.u., arbitrary units. Scale bars 10 μm.

Interestingly, the *cone-1/ceh-44* locus contains within its second and third intron predicted CUT homeodomain binding sites (DNA-binding sites of CUX and ONECUT proteins (Jolma et al. 2013)) which are typically present in pan-neuronal genes (Leyva-Diaz and Hobert 2022). Indeed, chromatin immunoprecipitation (ChIP) analysis conducted by the modENCODE consortium (Davis et al. 2018) demonstrated binding of the CUT factors CEH-48 and CEH-38 to these two different intronic regions. We find that transgenic reporters containing either of these putative binding sites drive robust neuronal expression (**Fig. 4A**). Employing CRISPR/Cas9 technology, we deleted the CUT homeodomain binding site within intron 3 in the *ceh-44/CUX* GFP-tagged endogenous locus, resulting in a significant reduction in *ceh-44/CUX* expression (**Fig. 4B**). Such reduction of *ceh-44::gfp* expression is also observed upon genetic removal of the five neuronally expressed ONECUT genes (*ceh-38*, *ceh-48*, *ceh-39*, *ceh-21* and *ceh-41*), which together with *ceh-44/CUX*, redundantly control pan-neuronal gene expression (Leyva-Diaz and Hobert 2022)(**Fig.3C**).

We next asked whether the CUT regulation of the *ceh-44/CUX* locus may be a reflection of CUT genes maintaining proper *ceh-44::gfp* expression after the initial onset of *ceh-44/CUX* expression. Indeed, we find that while expression of *ceh-44/CUX* is significantly reduced throughout all larval stages in quintuple CUT mutant animals, expression remained unaffected at the embryonic comma stage, during the onset of *ceh-44/CUX* expression (**Fig. 4C**). Therefore, we conclude that the CUT genes are required for the maintenance of *ceh-44/CUX* expression through binding to the intronic elements present in the *cone-1/ceh-44* locus. We surmise that this autoregulation is what produces the shorter *ceh-44/CUX* isoform that is observed in RNA-seq data in the nervous system (**Fig. 3A**) and that we mapped via GFP insertions into the locus (**Fig. 3F**).

### UNC-75/CELF is required for pan-neuronal *ceh-44/CUX* transcript production

While we have gained insights into the transcriptional regulation of the *cone-1/ceh-44* locus, unraveling the mechanisms underlying the alternative splicing of *ceh-44/CUX* and *cone-1/CASP* is crucial for understanding the generation of their distinct expression patterns. We initially focused this analysis on the two introns that precede the first isoform-specific exons of *ceh-44/CUX* and *cone-1/CASP*, respectively, namely intron 7 and intron 11 (**Fig.5A**), asking whether these introns may contain splicing regulatory sequences. To this end we developed a two-color splicing reporter cassette that provides a fluorescent readout of splicing regulation. The reporter cassette is expressed under control of a heterologous ubiquitous promoter and places GFP and RFP behind each of these introns, in lieu of the first isoform-specific exons of *ceh-44/CUX* and *cone-1/CASP* (**Fig.5A**). Hence, tissue-specific alternative splicing should become evident through tissue-specific expression of GFP and RFP. Mirroring the expression pattern of the endogenous proteins, we found that GFP expression from this reporter cassette is indeed detected in all neuronal cells, whereas RFP is detected in non-neuronal cell types (**Fig. 5A**).

**Figure 5:**
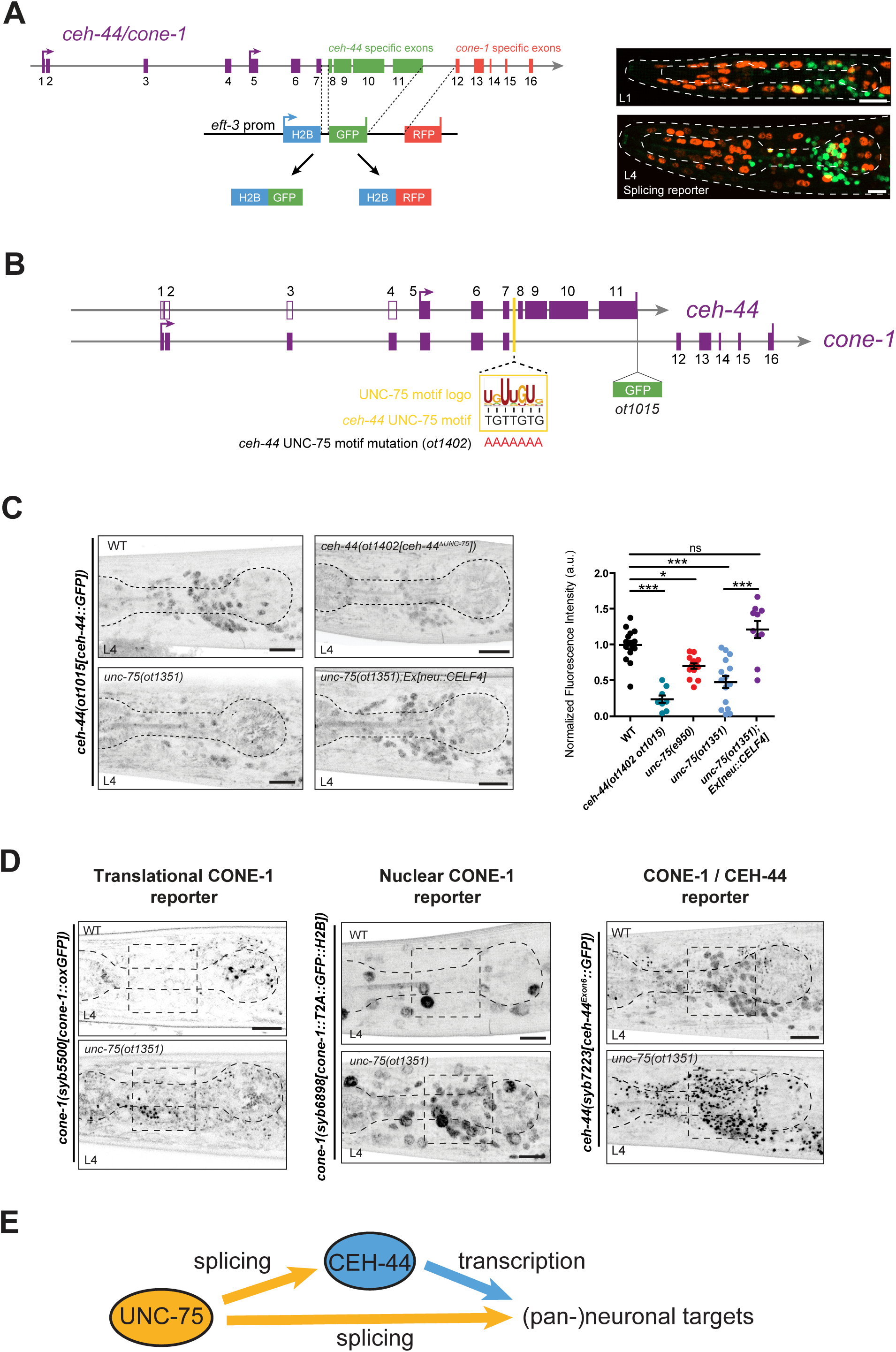
The pan-neuronal CELF homolog *unc-75/CELF* is required for *ceh-44/CUX* production. (**A**) Schematic representation of the splicing reporter cassette. GFP or RFP fluorescent proteins are localized to the nucleus via H2B and capture the alternative splicing activity of the gene locus. Images show lateral views in all panels (L4 stage in **A** and **B**). Images show lateral head views at the L1 (top) and L4 (bottom) larval stages (max Z-projections). Although sporadic instances of both reporters being expressed in the same cells are observed, these events are inconsistent, while the general splicing pattern remains reproducible and invariant from one animal to the next. (**B**) Schematic representation of *cone-1/ceh-44* gene locus showing GFP insertion, the UNC-75/CELF motif logo, location within intron 7, and motif mutation in *ceh-44(ot1402 ot1015)*. (**C**) *ceh-44(ot1015[ceh-44::gfp])* reporter expression in wild-type (top left), *ceh-44(ot1402 ot1015)* (top right), *unc-75(ot1351)* (bottom left), and *unc-75(ot1351);neu::CELF4[otEx8142]* (bottom right). Quantification of CRISPR allele fluorescence intensity in head neurons. Each dot represents the expression level within one worm with the mean and ± SEM indicated. Wild-type data are represented with black dots, *ceh-44(ot1402 ot1015)* with teal, *unc-75*(*e950*) with red, *unc-75(ot1351)* in blue, and *unc-75(ot1351);neu::CELF4[otEx8142]* in purple. One-way ANOVA followed by Tukey’s multiple comparisons test; *P < 0.05, ***P < 0.001. n ≥ 10 for all genotypes. (**D**) *cone-1(syb5500[cone-1::oxGFP])* (left), *cone-1(syb6898[cone-1::T2A::gfp::H2B])* (middle) and *ceh-44(syb7223[ceh-44Exon6::gfp])* (right) reporter expression in wild-type (top) and *unc-75(ot1351)* mutant animals (head, lateral view, L4 stage). Dashed square outlines the nerve ring region. (**E**) Model illustrating the feedforward architecture to control the production of pan-neuronal targets through UNC-75/CELF splicing and CEH-44/CUX transcriptional regulation. ns, not significant; a.u., arbitrary units. Scale bars 10 μm.

Since *cone-1/CASP* expression is ubiquitous at early embryonic stages and *ceh-44/CUX* expression becomes produced upon postmitotic neuronal differentiation, we hypothesized that a neuronal-specific splicing factor might be involved in the generation of neuron-specific *ceh-44/CUX* and the active exclusion of *cone-1/CASP*. The *C. elegans* homolog of the CELF family of RRM domain containing splicing factors, UNC-75/CELF, constitutes a pan-neuronally expressed splicing factor (Kuroyanagi et al. 2013a; Kuroyanagi et al. 2013b; Norris et al. 2014; Chen et al. 2016; Koterniak et al. 2020). Expression of *unc-75/CELF* was previously observed in all neurons through transgenic reporter constructs (Loria et al. 2003) and we sought to confirm this expression pattern with a CRISPR/Cas9 engineered endogenous reporter allele. We observed expression of this reporter allele in all neurons, starting at the embryonic comma stage, coinciding with the onset of *ceh-44/CUX* expression (**Fig. S4A**). *unc-75/CELF* pan-neuronal expression is maintained through all larval stages and into adulthood, aligning with the expression pattern of *ceh-44/CUX* and positioning *unc-75/CELF* as a strong candidate to regulate the *cone-1/ceh-44* splicing event. Supporting this notion, animals carrying the canonical *unc-75(e950)* mutant allele, a deletion affecting two of the three UNC-75/CELF RNA binding RRM motifs (Loria et al. 2003) (**Fig. S4A**) show a strong reduction in *ceh-44/CUX* expression (**Fig. 5C**). We also engineered a more unambiguous null allele, *unc-75(ot1351)*, in which all three RRMs are removed (**Fig. S4A**). *unc-75(ot1351)* animals display locomotory defects similar to those observed in *unc-75(e950)* (**Fig. S4B**), and expression of *gfp-*tagged *ceh-44/CUX* was also significantly reduced (**Fig. 5C**).

UNC-75/CELF regulation of *ceh-44/CUX* splicing is likely direct. A predicted UNC-75/CELF binding site (Koterniak et al. 2020) is located within intron 7 of the *cone-1/ceh-44* locus, between the shared exons and *ceh-44/CUX* and *cone-1/CASP* specific exons (**Fig. 5B**). Furthermore, an UNC-75/CELF binding peak was observed in this intron through CLIP-seq analysis (John Calarco, pers. comm.). We assessed the functional relevance of this UNC-75/CELF binding site through deletion from the endogenous *cone-1/ceh-44* locus (**Fig. 5B**, *ot1402* allele). Mutation of the UNC-75/CELF binding site significantly reduced *ceh-44/CUX* expression, as visualized with the endogenous *ceh-44/CUX* CRISPR reporter (**Fig. 5C**).

Consistent with this loss of *ceh-44/CUX* expression, transcription of the pan-neuronal gene *ric-19* is significantly reduced in *unc-75/CELF* mutants, indicating defective *ceh-44/CUX* function (**Fig. S4C**). Synaptic transmission defects in *unc-75/CELF* mutants can be rescued by expressing a human *CELF4* cDNA under the control of the pan-neuronal *unc-75/CELF* promoter (Loria et al. 2003). Here we demonstrate that the same construct can rescue the *ceh-44/CUX* expression phenotype of *unc-75/CELF* mutants (**Fig. 5C**), confirming the conservation of UNC-75/CELF function across phylogeny.

Given these findings, we reasoned that if UNC-75/CELF indeed controls a neuron-specific splicing event that promotes *ceh-44/CUX* transcript production at the expense of *cone-1/CASP*, *unc-75/CELF* mutant animals should exhibit ectopic neuronal *cone-1/CASP* expression. We observed such ectopic expression using three different *cone-1/CASP* reporter alleles (**Fig. 5D**). Altogether, the neuronal expression of the different *cone-1/CASP* reporter alleles in *unc-75/CELF* mutant animals indicate a mechanism through which UNC-75/CELF promotes *ceh-44/CUX* transcript production in the nervous system while excluding the alternative, *cone-1/CASP-*encoding transcript, from the nervous system.

## DISCUSSION

In this study, through extensive use of CRISPR/Cas9 technology, we have deconstructed the architecture and regulation of a phylogenetically deeply conserved and highly unusual genetic locus that produces two distinct proteins with no apparent functional overlap. We propose the following model for *cone-1/ceh-44* transcriptional regulation (**Fig. S5A**): transcript production from this locus is directed to all cells by factors controlling the upstream promoter, which is active throughout the life of the organism. Upon neuronal differentiation, the onset of UNC-75/CELF locus switches transcript production in neurons to *ceh-44/CUX*, on the expense of *cone-1/CASP*. In neurons, *ceh-44/CUX* expression is then maintained by CUT factors, including CEH-44/CUX, binding to elements upstream of the *ceh-44/CUX* transcript start.

UNC-75/CELF, a conserved RNA-binding protein, plays a crucial role in the regulation of RNA metabolism, including the regulation of splicing (Dasgupta and Ladd 2012; Kuroyanagi et al. 2013a; Kuroyanagi et al. 2013b; Norris et al. 2014; Chen et al. 2016; Koterniak et al. 2020). In *C. elegans*, UNC-75, along with other splicing factors, orchestrates splicing networks that contribute to neuronal diversity (Norris et al. 2014; Chen et al. 2016; Koterniak et al. 2020). Interestingly, almost half (235) of the 534 genes bound by UNC-75 in CLIP-Seq analysis (Chen et al. 2016) are also bound in CHIP-Seq analysis of several CUT homeodomain proteins (Davis et al. 2018). Hence, through directing CEH-44/CUX CUT homeodomain expression to the nervous system through alternative splicing, UNC-75/CELF orchestrates in a feedforward manner the production of CEH-44/CUX-dependent primary transcripts that then serve as its own substrates for splicing (**Fig.5E**).

Our findings show that *cone-1/CASP* is actively excluded from the nervous system, and we have also identified another golgin with a cell type-specific expression pattern. This contrasts with most golgins, large ubiquitously expressed coiled-coil proteins that decorate the cytoplasmic surface of the Golgi apparatus, extending tentacle-like structures into the cytosol (Munro 2011; Gillingham and Munro 2016; Lowe 2019). They are thought to not only capture incoming vesicles but also provide specificity to such tethering step (Munro 2011; Gillingham and Munro 2016; Lowe 2019). The identification of two Golgi proteins with cell-type specific expression patterns points to cell-type specific functions of the Golgi apparatus.

Notably, *cone-1/ceh-44* locus structure is highly conserved throughout evolution. Early studies on CASP expression indicated that the transcript of this isoform is widely distributed througout mouse tissues (Lievens et al. 1997), similar to *cone-1/CASP* in *C. elegans*. Recent studies using CASP isoform specific antibodies have found that CASP is also widely expressed in neurons across multiple brain regions (Suzuki et al. 2022). *Cux1* is present in various brain regions throughout development and in adult neurons but is also found in multiple non-neuronal tissues, unlike *C. elegans ceh-44/CUX* (Rong Zeng et al. 2000; Weiss and Nieto 2019). Therefore, while *Cux1* locus structure resembles that of *cone-1/ceh-44*, the expression patterns of each transcript differ through phylogeny. Still, expression pattern analysis with single cell resolution may reveal mutually exclusive sites of expression. Functionally, although we did not find any phenotype in *cone-1/CASP* null animals, a recent study has identified CASP variants in patients with temporal lobe epilepsy (Suzuki et al. 2022). Analysis of *Casp*-specific mutant mice indicated that these variants affect neuronal properties and enhance excitatory synaptic transmission. Another study found a higher proportion of motor phenotypes in individuals with variants affecting both CUX1 and CASP, compared to those with variants affecting only CUX1 protein (Oppermann et al. 2023). Future studies are needed to clarify the role of CASP variants, and to understand the spatial and temporal production control of CUX1/CASP, including the potential involvement of the homologs of UNC-75, the CELF proteins.

## MATERIALS AND METHODS

### Caenorhabditis elegans strains and handling

Worms were grown at 20°C on nematode growth media (NGM) plates seeded with *E. coli* (OP50) bacteria as a food source (Brenner 1974). Wild-type strain used is Bristol variety, strain N2. A complete list of strains used in this study is listed in **Table S1**. Information about transgenes and alleles generated via CRISPR/Cas9-based genome engineering is detailed in Supplemental Materials and Methods.

### Automated worm tracking

For the tracking of *unc-75/CELF* mutants and controls, tracking was performed using the WormLab automated multiworm tracking system (MBF Bio-science) (Roussel et al. 2014) at room temperature. In each plate, 5 young adult animals were recorded for 5 minutes and tracked on NGM plates with no food. Videos were segmented to extract the worm contour and skeleton for phenotypic analysis. Raw WormLab data was exported to Prism (GraphPad) for further statistical analysis. Statistical significance between each group was calculated using One-way ANOVA followed by Tukey’s multiple comparison’s test.

### Microscopy

Worms were anesthetized using 100 mM of sodium azide and mounted on 5% agarose on glass slides. Images were acquired using a Zeiss confocal microscope (LSM 980) and a Zeiss fluorescence microscope (Axio Imager.Z1). Image reconstructions were performed using Zeiss Zen software tools. Maximum intensity projections of representative images are shown. Fluorescence intensity was quantified using the ImageJ software (Schneider et al. 2012). Figures were prepared using Adobe Illustrator.

### Colocalization analysis

Colocalization analysis were performed by preparing the fluorescence microscopy images with ImageJ software (Schneider et al. 2012) and analyzing them with the JACoP plugin. Their relative degree of colocalization was expressed in Pearson’s Correlation Coefficient (Adler and Parmryd 2010).

### Visualization of averaged RNA-Seq coverage tracks

Alignment files from our previous nervous system transcriptional profiling (Leyva-Diaz and Hobert 2022) (GEO accession #GSE188489) were used to generate an averaged coverage track with bedtools and then visualized with the Integrative Genomics Viewer (IGV) genome browser (Robinson et al. 2011).

### Quantification and statistical analysis

All microscopy fluorescence quantifications were done in the ImageJ software (Schneider et al. 2012). For all images used for fluorescence intensity quantification, the acquisition parameters were maintained constant among all samples (same pixel size, laser intensity), with control and experimental conditions imaged in the same imaging session. For quantification of head neurons, fluorescence intensity was measured in maximum intensity projections using a single rectangular region of interest from anterior to posterior bulb. A standard threshold was assigned to all the control and experimental conditions being compared. All statistical tests for fluorescence quantifications and behavior assays were conducted using Prism (Graphpad) as described in figure legends.

## ACKNOWLEDGEMENTS

We thank Chi Chen for generating transgenic strains, Barth Grant for sharing the *aman-2* reporter (RT2745 strain), HaoSheng Sun, Nuria Flames and members of the Hobert lab and Leyva-Díaz lab for commenting on the manuscript, WormBase (Sternberg et al. 2024) and CGC (funded by NIH Office of Research Infrastructure Programs, P40 OD010440) for providing resources and reagents. This work was funded by the Howard Hughes Medical Institute (O.H.) and by a CIDEGENT grant from Generalitat Valenciana (CIDEXG/2022/30) (E.L.-D.).

## AUTHOR CONTRIBUTIONS

E.L-D. and O.H. conceived the project and designed the experiments. E.L-D. and M.C. performed all the experiments, imaging and quantifications, except for those specified. K.P. conducted the experiments, imaging and quantifications shown in **Fig. 4B** and **S4A**. J.I.J.-L. and J.V. performed the quantifications shown in **Fig. 1D** and **2E**. The manuscript was prepared by E.L-D., M.C. and O.H.

