## Supplemental Material for "Alternative splicing controls pan-neuronal homeobox gene expression"

**Supplemental Materials and Methods** includes detailed information about transgenes and CRISPR allele generation.

**Supplemental Table S1** includes all the *C. elegans* strains used in this study.

**Supplemental Figure S1** related to Figure 2, shows a characterization of *cone-1/CASP* mutants.

**Supplemental Figure S2** related to Figure 3, shows CEH-44/CUX protein structure and sequence details.

**Supplemental Figure S3** related to Figure 4, shows the expression analysis of an endogenous *cone-1/CASP* transcriptional reporter.

**Supplemental Figure S4** related to Figure 5, shows the characterization of new *unc-75/CELF* reporter and null alleles.

**Supplemental Figure S5** related to discussion, shows a model for *ceh-44/CUX* and *cone-1/CASP* transcript production.

### SUPPLEMENTAL MATERIALS AND METHODS

#### CRISPR/Cas9-based genome engineering

*cone-1(ot1485 syb5500), ceh-44(ot1486 ot1015)*, *ceh-44(ot1529 ot1015), unc-75(ot1351 ot1015), cone-1(syb5437 ot1410), ceh-44(ot1447 ot1015), ceh-44(ot1402 ot1015), cone-1(ot1502)* were generated using Cas9 protein, tracrRNA, and crRNAs from IDT, as previously described (Dokshin et al. 2018). For *cone-1(ot1485 syb5500)* and *ceh-44(ot1486 ot1015)*, one crRNA (agtttgacggtcttctagca) and a ssODN donor (tgatcaaggcatttcaaagcgagtttgacgggtctttaagcacggagcacagctgctgaaaatgcactgattga) were used to introduce a nonsense mutation in *cone-1/ceh-44* exon 3. For c*eh-44(ot1529 ot1015),* two crRNAs (agcagtattggcgggacgcg and ctaatacggacgggtgacgg) and a ssODN (cttctcatgggggttgttggtgcctcccgccccgccggcggctctttttgtgctcaatttttcttgatc) were used to delete 224 bp overlapping with the CEH-38/48 ChIP peak on *ceh-44*. For *unc-75(ot1351 ot1015)* two crRNAs (ttaaaacggttcgaagttgg and taatcaatcaataatggcac) and a ssODN as a donor (ttaatggcctaacattttgtatttctaggccaccaccattattgattgattatatatgtatttttgtatt) were used to generate a 7665 bp deletion in *unc-75*. For *cone-1(syb5437 ot1410)* and *ceh-44(ot1447 ot1015)*, two crRNAs (cttttttcgacttcgttga and gctaaacacaaaattgtatg) and a ssODN (caagattttcaatttttcagatcgaaaacgccgtcaagcataagattgtatgcggtttactgagagaatttacatattcccga) were used to delete *cone-1/ceh-44* exon 5. For UNC-75 binding site mutation in *ceh-44(ot1402 ot1015)*, the UNC-75 motif UGUUGUG (+8992, +8998) was mutated to AAAAAAA. For *cone-1(ot1502[GFP::H2B::SL2::cone-1]),* a crRNA (ctcttgagacgatttccata) and a ssODN
(ttaaataattcgattaatttcttatttcagaccttatgagtaaaggagaagaacttttcactggagttgtcccaattcttgttgaatt
agatggtgatgttaatgggcacaaattttctgtcagtggagagggtgaaggtgatgcaacatacggaaaacttacccttaa
atttatttgcactactggaaaactacctgttccatgggtaagtttaaacatatatatactaactaaccctgattatttaaattttcag
ccaacacttgtcactactttctgttatggtgttcaatgcttctcgagatacccagatcatatgaaacggcatgactttttcaagag
tgccatgcccgaaggttatgtacaggaaagaactatatttttcaaagatgacgggaactacaagacacgtaagtttaaaca
gttcggtactaactaaccatacatatttaaattttcaggtgctgaagtcaagtttgaaggtgatacccttgttaatagaatcgag
ttaaaaggtattgattttaaagaagatggaaacattcttggacacaaattggaatacaactataactcacacaatgtatacat
catggcagacaaacaaaagaatggaatcaaagttgtaagtttaaacatgattttactaactaactaatctgatttaaattttca
gaacttcaaaattagacacaacattgaagatggaagcgttcaactagcagaccattatcaacaaaatactccaattggcg

atggccctgtccttttaccagacaaccattacctgtccacacaatctgccctttcgaaagatcccaacgaaaagagagacc acatggtccttcttgagtttgtaacagctgctgggattacacatggcatggatgaactatacaaaccaccaaagccatctgc caagggagccaagaaggccgccaagaccgttacgaagccaaaggacggaaagaagagacgtcatgcccgtaagg aatcatactccgtctacatctaccgtgtcctcaagcaagttcatccagacactggagtttcctccaaagccatgtctatcatga actcttttgtcaacgatgtcttcgagcgtattgctgctgaagcatcccgtcttgctcactacaacaagcgttccacaatctcatc ccgcgaaattcagaccgctgtccgtctgatccttccaggagagcttgccaagcacgccgtgtctgagggaaccaaggcc gttaccaagtacacttccagcaagtaggctgtctcatcctactttcacctagttaactgcttgtcttaaaatctatgcttctctttag tatctaaaattttcctagaagcttacaagtatataaatggtctcttctcaataaaggttgtatatttattcatcttattgaatctgccat ttcctcgtttttgcgagtttatataccttccaattttctttctattgtattttcaacttctaattttaattcagggaaactgcttcaacgcat catggaaatcgtctcaagagcatgggaatctgtagatt) were used to insert an H2B::SL2 sequence in *cone-1(syb5437[GFP::cone-1])* between the GFP reporter and the gene.
*cone-1(syb5500)*, *cone-1(syb6898)*, *golg-2(syb6680)*, *golg-4(syb6547)*, *golg-5(syb5400)*, *cone-1(syb5437)*, *cone-1(syb7529)*, *ceh-44(syb7223)*, *ceh-44(syb6281)*, *ceh-44(syb5843)*, *unc-75(syb6499)* were generated by SUNY Biotech.

**Reporter transgenes**

The *ceh-44 cis*-regulatory element reporters were generated using a PCR fusion approach (Hobert 2002). The *ceh-44prom1* (-2000, -1), *ceh-44prom2* (-26, -1), *ceh-44prom3* (+2075, +2560), *ceh-44prom4* (+4708, +4980), *ceh-44prom5* (+3330, +6689) promoter fragments were amplified from N2 genomic DNA and and fused to *2xNLS-GFP*. *ceh-44prom3* and *ceh-44prom4* promoter fragment coordinates match those of the CEH-38/48 ChIP peaks in the regulatory regions of this gene. *ceh-44prom5* contains the entirety of intron 3, exon4, and intron 4 through the last possible in frame start codon on exon 5. The resulting PCR fusion DNA fragments were injected as simple extrachromosomal arrays (50 ng/mL) into *pha-1(e2123)* animals, using a *pha-1* rescuing plasmid (pBX at 50 ng/µL) as co-injection marker. Extrachromosomal array lines were selected according to standard protocol.

**Table S1. Strains used in this study.** Promoter coordinates in relation to the ATG.

| Strain name | Strain genotype | Source |
|---|---|---|
| N2 | *C. elegans* Strain N2 (WormBase: WBStrain00000001) | CGC |
| CB950 | *unc-75(e950)* | (Loria et al. 2003) |
| OH10690 | *otIs356(rab-3prom1::2xNLS-tagRFP) V* | (Stefanakis et al. 2015) |
| OH11062 | *otIs381(ric-19prom6::2xNLS-GFP) V* | (Stefanakis et al. 2015) |
| OH16219 | *ceh-44(ot1015[ceh-44::gfp]) III* | (Leyva-Diaz and Hobert 2022) |
| OH16377 | *ceh-38(tm321) II; ceh-44(ot1028) III; ceh-48(tm6112) IV; otIs356 V; otDf1 X* | (Leyva-Diaz and Hobert 2022) |
| OH18168 | *ceh-44(ot1015[ceh-44::gfp]), cone-1(ot1282) III* | This study |
| OH18267 | *cone-1(syb5500[cone-1::oxGFP]), golg-4(syb6547[wrmScarlet::golg-4]) III; him-5(e1490) V* | This study |
| OH18268 | *ceh-44(ot1402[ceh-44*$^{\Delta UNC-75}$*]) ot1015[ceh-44::gfp]) III* | This study |
| OH18318 | *unc-75(ot1351) I; otIs381(ric-19prom6::2xNLS-GFP) V* | This study |
| OH18319 | *cone-1(ot1287) III; otIs381(ric-19prom6::2xNLS-GFP) V* | This study |
| OH18347 | *golg-2(syb6680[wrmScarlet::golg-2]) II; cone-1(syb5500[cone-1::oxGFP]) III* | This study |
| OH18418 | *unc-75(e950) I; ceh-44(ot1015[ceh-44::gfp]) III* | This study |
| OH18426 | *otEx8048(ceh-44prom1::2xNLS-GFP, pha-1(+)); pha-1(e2123) III* | This study |
| OH18429 | *otEx8050(ceh-44prom3::2xNLS-GFP, pha-1(+)); pha-1(e2123) III* | This study |
| OH18463 | *unc-75(ot1351) I; ceh-44(ot1015[ceh-44::gfp]) III* | This study |
| OH18465 | *ceh-38(tm321) II; ceh-44(ot1015[ceh-44::gfp]) III; ceh-48(tm6112) IV; otIs356 V; otDf1 X* | This study |
| OH18495 | *otEx8077(ceh-44prom4::2xNLS-GFP, pha-1(+)); pha-1(e2123) III* | This study |
| OH18498 | *otEx8080(ceh-44prom2::2xNLS-GFP, pha-1(+)); pha-1(e2123) III* | This study |
| OH18577 | *cone-1(syb5500[cone-1::oxGFP]) III; pwIs1022[snx-1prom::aman-2::tagRFP]* | This study |
| OH18750 | *cone-1(syb5437[gfp::cone-1] ot1410[cone-1*$^{\Delta Exon5}$*]) III* | This study |
| OH18751 | *unc-75(ot1351) I; cone-1(syb5500[cone-1::oxGFP]) III* | This study |
| OH18752 | *unc-75(ot1351) I; cone-1(syb6898[cone-1::T2A::gfp::H2B])* | This study |
| OH18790 | *otEx8142(unc-75prom1::CELF4, inx-6prom::tagRFP); unc-75(ot1351) I; ceh-44(ot1015[ceh-44::gfp]) III* | This study |
| OH18841 | *unc-75(ot1351) I; ceh-44(syb7223[ceh-44*$^{Exon6}$*::gfp]) I* | This study |
| OH18948 | *ceh-44(ot1447[ceh-44*$^{\Delta Exon5}$*] ot1015[ceh-44::gfp]) III* | This study |
| OH18958 | *otEx8112(eft-3prom::H2B::ceh-44intron7::gfp::ceh-44intron11::tagRFP)* | This study |
| OH19071 | *otEx8213(ceh-48prom4::ceh-44(E8-11)); ceh-38(tm321) II; ceh-44(ot1028) III; ceh-48(tm6112) IV; otIs356 V; otDf1 X* | This study |
| OH19076 | *otEx8218(ceh-44prom5::2xNLS-GFP, pha-1(+)); pha-1(e2123) III* | This study |
| OH19119 | *cone-1(ot1485[Exon3STOP] syb5500[cone-1::oxGFP]) III* | This study |
| OH19186 | *cone-1(ot1502[gfp::H2B::SL2::cone-1]) III* | This study |
| OH19210 | *ceh-44(ot1486[Exon3STOP] ot1015[ceh-44::gfp]) III* | This study |
| OH19271 | *ceh-44(ot1529[ceh-44*$^{\Delta CUT}$*]) ot1015[ceh-44::gfp]) III* | This study |
| PHX5400 | *golg-5(syb5400[golg-5::wrmScarlet]) I* | This study |
| PHX5437 | *cone-1(syb5437[gfp::cone-1]) III* | This study |
| PHX5500 | *cone-1(syb5500[cone-1::oxGFP]) III* | This study |

| | | |
|---|---|---|
| PHX5843 | *ceh-44(syb5843[ceh-44$^{Exon8}$::gfp]) III* | This study |
| PHX6281 | *ceh-44(syb6281[ceh-44$^{Exon7}$::gfp]) III* | This study |
| PHX6499 | *unc-75(syb6499[gfp::unc-75]) I* | This study |
| PHX6547 | *golg-4(syb6547[wrmScarlet::golg-4]) III* | This study |
| PHX6680 | *golg-2(syb6680[wrmScarlet::golg-2]) II* | This study |
| PHX6898 | *cone-1(syb6898[cone-1::T2A::gfp::H2B]) III* | This study |
| PHX7223 | *ceh-44(syb7223[ceh-44$^{Exon6}$::gfp]) III* | This study |
| PHX7529 | *cone-1(syb7529[cone-1$^{Exon4}$::gfp]) III* | This study |

**Supplemental Figure 1: *ceh-44/CUX* does not require *cone-1/CASP* for expression or function**

(**A**) Schematic representation of the *cone-1/ceh-44* gene locus showing mutant alleles (red asterisks indicate early stop codons).

(**B-C**) *ceh-44(ot1015[ceh-44::gfp])* (**B**) and *ric19prom6::2xNLS-GFP[otIs381]* (**C**) reporter expression in L4 animals (head, lateral views) in wild-type (top) and *cone-1/CASP* mutants (bottom). Note that the *ot1282* (**B**) and *ot1287* (**C**) mutant alleles are designed to introduce a frameshift and harbor the same molecular lesion. Quantification of fluorescence intensity in head neurons. The data are presented as individual values with each dot representing the expression level of one worm with the mean ± SEM indicated. Unpaired *t*-test, n ≥ 10 for all genotypes.

ns, not significant; a.u., arbitrary units. Scale bars 10 μm.

**Supplemental Figure 2: GFP insertions throughout CONE-1/CEH-44 sequence**

(**A**) AlphaFold (Jumper et al. 2021) prediction of CEH-44/CUX protein structure with GFP insertion locations.

(**B**) CEH-44/CUX protein sequence (Exon 1-11). Residues coded in odd exons are shown in bold, GFP insertion locations depicted in **S3A** are denoted with a caret (^) and highlighted in purple, *ceh-44/CUX* specific exons (8-11) are highlighted in grey, methionines within the shared exons (1-7) are shown in red.

(**C**) *ceh-44/CUX* exon 5 DNA sequence. Deleted region in *ot1410* and *ot1447* alleles (**Fig. 3A, D, E**) is underlined, and the preserved sequence (in frame) is shown in bold. Three in frame initiation codons are highlighted in yellow.

**Supplemental Figure 3: Endogenous *cone-1/ceh-44* transcriptional reporter expression**

(**A**) Schematic representation of the *cone-1/ceh-44* gene locus showing the *gfp::H2B::SL2* cassette insertion location.

(**B**) Temporal expression analysis in *cone-1(ot1502[gfp::H2B::SL2::cone-1])* across different embryonic stages (2-Cell, 44-Cell, Comma, 1.5-Fold, 2-Fold) and larval stages (L1 and L4). All images show lateral view.

Scale bars 10 μm.

**Supplemental Figure 4: Characterization of *unc-75/CELF* expression and function with novel reporter and null alleles.**

(A) Schematic representation of the *unc-75/CELF* gene locus showing the location of the RNA recognition motifs (RRM), GFP insertion and mutant deletion alleles (classic *e950* and *ot1351*, generated in this study). *unc-75(syb6499[gfp::unc-75])* reporter expression shown at the comma embryonic stage (bottom left, lateral view), L1 larval stage (top, full worm lateral view) and young adult stage (bottom right, lateral view of the head). The embryonic comma stage is the stage when neurons are born. Head ganglia, ventral nerve cord, and tail ganglia outlined in L1 image, and head ganglia outlined in young adult image. Asterisk (*) indicate autofluorescence in L1 gut.

(**B**) Worm locomotory features: center point speed (top left), reverse distance (top right), idle time (bottom left), and straight-line distance (bottom right). Each dot represents one worm with the mean ± SEM indicated. Wild-type data are represented with black dots, *unc-75(e950)* with red, and *unc-75(ot1351)* with teal. One-way ANOVA followed by Tukey's multiple comparisons test; ***p < 0.001. n ≥ 70 for all genotypes.

(**C**) *ric19prom6::2xNLS-GFP[otIs381]* reporter expression in L4 animals (head, lateral views) in wild-type (top) and *unc-75(ot1351)* (bottom) animals. The data are presented as individual values with each dot representing the expression level of one worm with the mean ± SEM indicated. Wild-type data are represented with black dots and *unc-75(ot1351)* mutants with red dots. Unpaired *t*-test, ***P < 0.001. n = 10 for all genotypes. YA, young adult; ns, not significant; a.u., arbitrary units. Scale bars 10 μm.

**Supplemental Figure 5: Model for *cone-1/ceh-44* transcript production**

(A) Model for *cone-1/ceh-44* production through transcriptional regulation and alternative splicing. At the DNA level, transcription is regulated by factors controlling the upstream promoter in all cells, and by CUT factors in neurons. In neurons, splicing of the *cone-1/ceh-44* pre-mRNA controlled by UNC-75/CELF leads to the production of *ceh-44/CUX*, while non-neuronal cells lacking UNC-75/CELF produce the *cone-1/CASP* transcript. Together with the other neuronally expressed CUT factors, CEH-44/CUX can control its own expression through autoregulation.

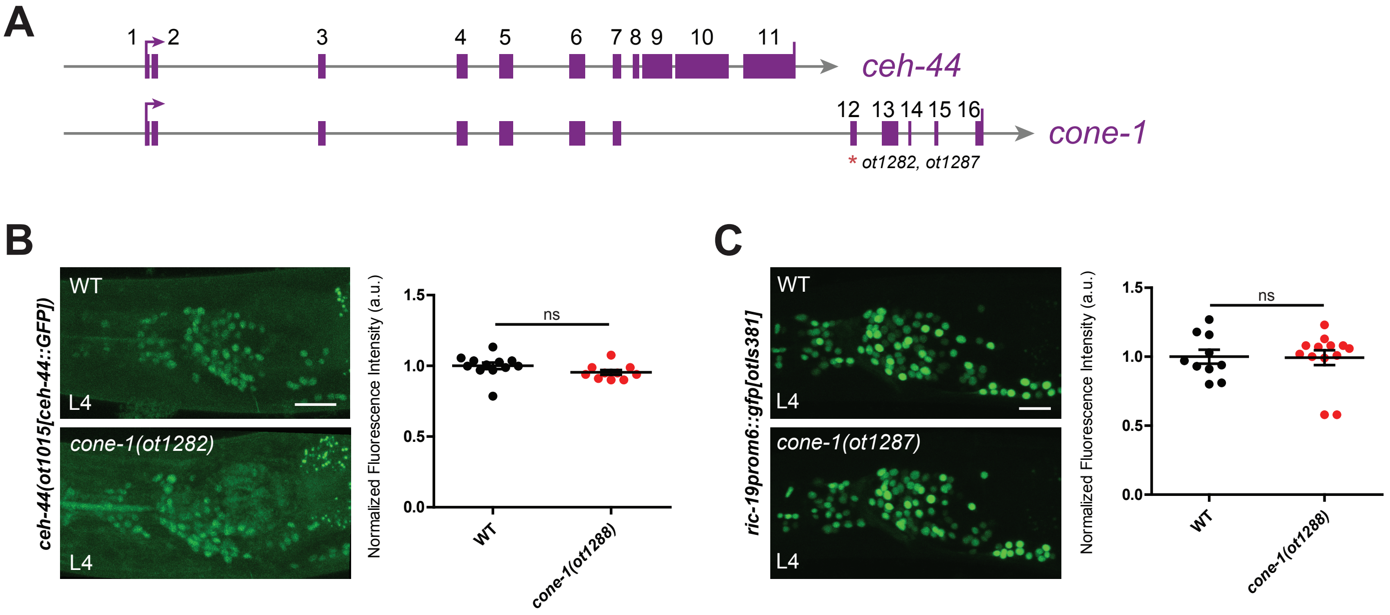
A
1
2
3
4
5
6
7
8
9
10
11
ceh-44
12
13
14
15
16
cone-1
* ot1282, ot1287
B
ceh-44(ot1015[ceh-44::GFP])
WT
L4
cone-1(ot1282)
L4
Normalized Fluorescence Intensity (a.u.)
1.5
1.0
0.5
0.0
ns
WT
cone-1(ot1288)
C
ric-19prom6::gfp[otIs381]
WT
L4
cone-1(ot1287)
L4
Normalized Fluorescence Intensity (a.u.)
1.5
1.0
0.5
0.0
ns
WT
cone-1(ot1288)

**A**

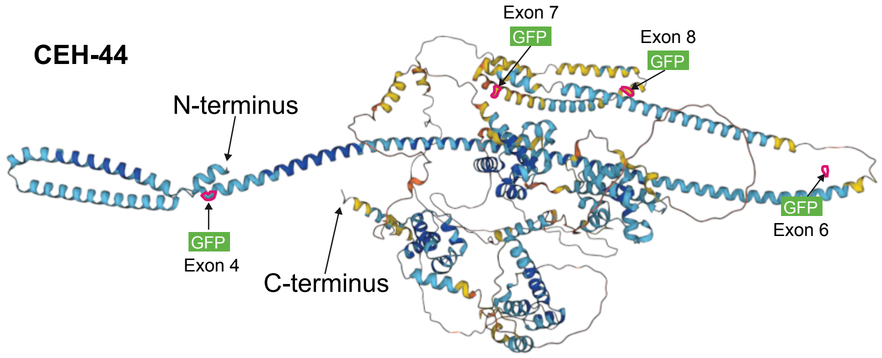

**B**

>CEH-44

**MEIVSRAWESVDWDRIQT**RVEAEVTALGQRQDDSEIRKTRLVEESNAYRGRTNK**DSRKVAIPLIKAFQSEFDGLLARSTAAENALIDICKSIVSLP**DPKS^LLKGAEAWKN
DAEKTQKAVEEREELKRQLIKVNNELEDLRGKDVKVRKLKDKLAKLESEQDIF**IENAVNEVEKKAEQELNDRLTELIAEKEKMKEQNEILEKNMDSLESKNKDIQRKLEIA**
**KQTVEQKDGLENEQLSIAMKDLADAKHKI**VFLEERVSQL^ENEAEKVNESKKAGNIEDIAALGSVLVQKDDVIQQLTNDIKRHEASHVEELAKWKLAVSAVEKKNKTLIGE
LNELKNQLESRNDYEAIKNELR**LLREIEFGDSAEANA^ESIERLGETVETLDRLLAEKNRRLQNENASLRVANDGFK**GDEVM^KAIVSGSHSRVVETVGKRVGAEEANSYR
QKNTDSELIEK**IQEAKRNKAVCELKFEDPTINVLTYLKNQKAKEAGKRDAPTPILAAPVTPKHVTKLGTHTITTTALPPRTQTAETTQSILQRLSNGSNKHLLNEDLKLST**
**VLNLKRFSGNPAKPLEAKTSEEQKAELETIEKMQKRIQVNVQALNGHPLNTTEIASHCKRLMIAYNIGQRLFAKHVMNQVVK**SQGSLSELLSKPRHWNKLTDKGREAFRRI
YGWISDDEAINLLCSLSPRRVWPADQNIEHPKAETLLDTSDPMEFKEEPVIRYDVTPKVEPVIEKIKSPVESPCSSQAGASSLRASRWRHDDISKEKILSILQTELKKIEE
ETTESKVVVPVKPTATGGNRRFSSNSTYESSSLTGKSRPSTVELILKQRISTGLQPLTQAQYDAYTVLDTDFLVKQIKEFLTMNSISQRQFGEYILGLSQGSVSDLLARPK
TWAQLTQKGREPFIRMQLFMDDVEASEENDEKQPKITICDEDSDLAKTLATLLNAVHREPSEPKTSVKLEPLSEIDV**IMQVPSASKPSSVVKSIDESSGEEILDTFEIVYQ**
**VKGILEENGISPRVFGDEYLHCTSSMCADLMIRTKSFENSKASEKLMYTRMKTFLSDPIAIPLLVEKEESKETVKAKIESVPAPREAPRPVKRKHSSDTDDYDLNTKKPIQ**
**RTVITDYQKDTLRFVFVNEQHPSNELCEQISLKLDMSLRTVQNWFHNHRTRSKAREKEGKVYSDALPNGTAVKSLTWKDDLQKMLDEAPAITSQWAPDYQNRAGSVKSSTS**
**ADSPTNNNYSSPIFSFDKASTTSTVKKPSSTGKLDNLVARMIRLAEGREAAAAKAS**

**C**

>*ceh-44* Exon 5

**ATCGAAAACGCCGTCA**ACGAAGTCGAAAAAAAGGCGGAACAAGAGCTTAACGATCGTCTGACAGAGTTAATCGCTGAAAAAGAGAAAATGAAAGAACAAAATGAGATTTTG
GAGAAGAATATGGATAGTCTCGAGTCGAAGAATAAGGATATTCAAAGAAAACTCGAAATTGCCAAACAAACTGTCGAGCAGAAGGATGGGCTCGAGAATGAGCAACTTTCG
ATTGCAATGAAGGATTTGGCTGATGCTA**AACACAAAATT**

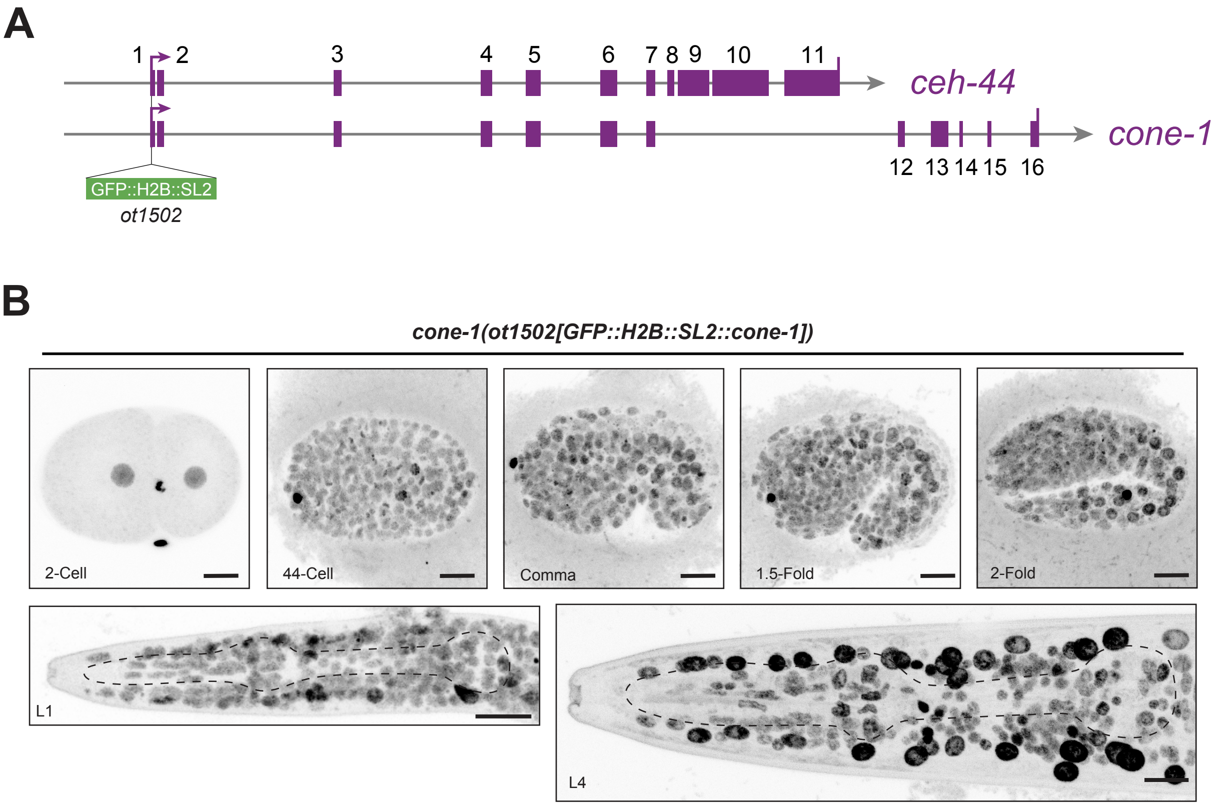
A
1
2
3
4
5
6
7
8
9
10
11
ceh-44
cone-1
12
13
14
15
16
GFP::H2B::SL2
ot1502
B
cone-1(ot1502[GFP::H2B::SL2::cone-1])
2-Cell
44-Cell
Comma
1.5-Fold
2-Fold
L1
L4

Leyva-Díaz_Figure_S4

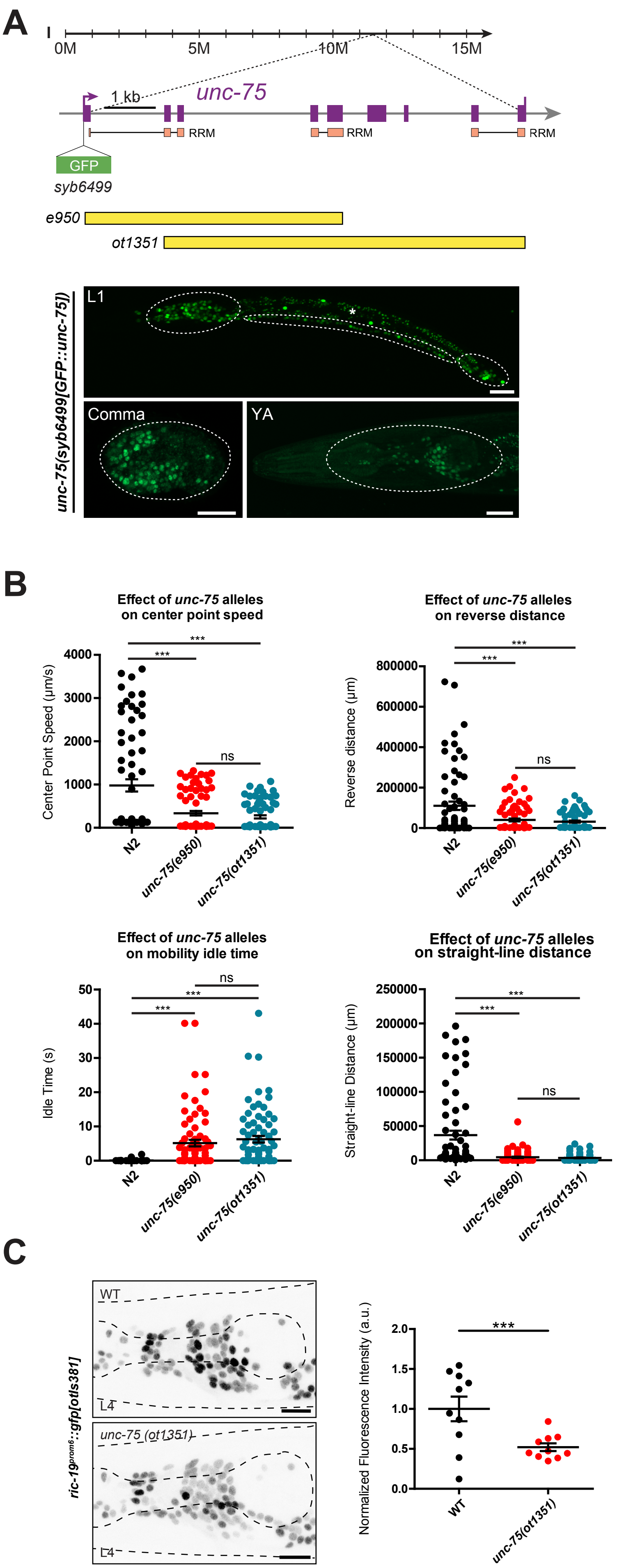

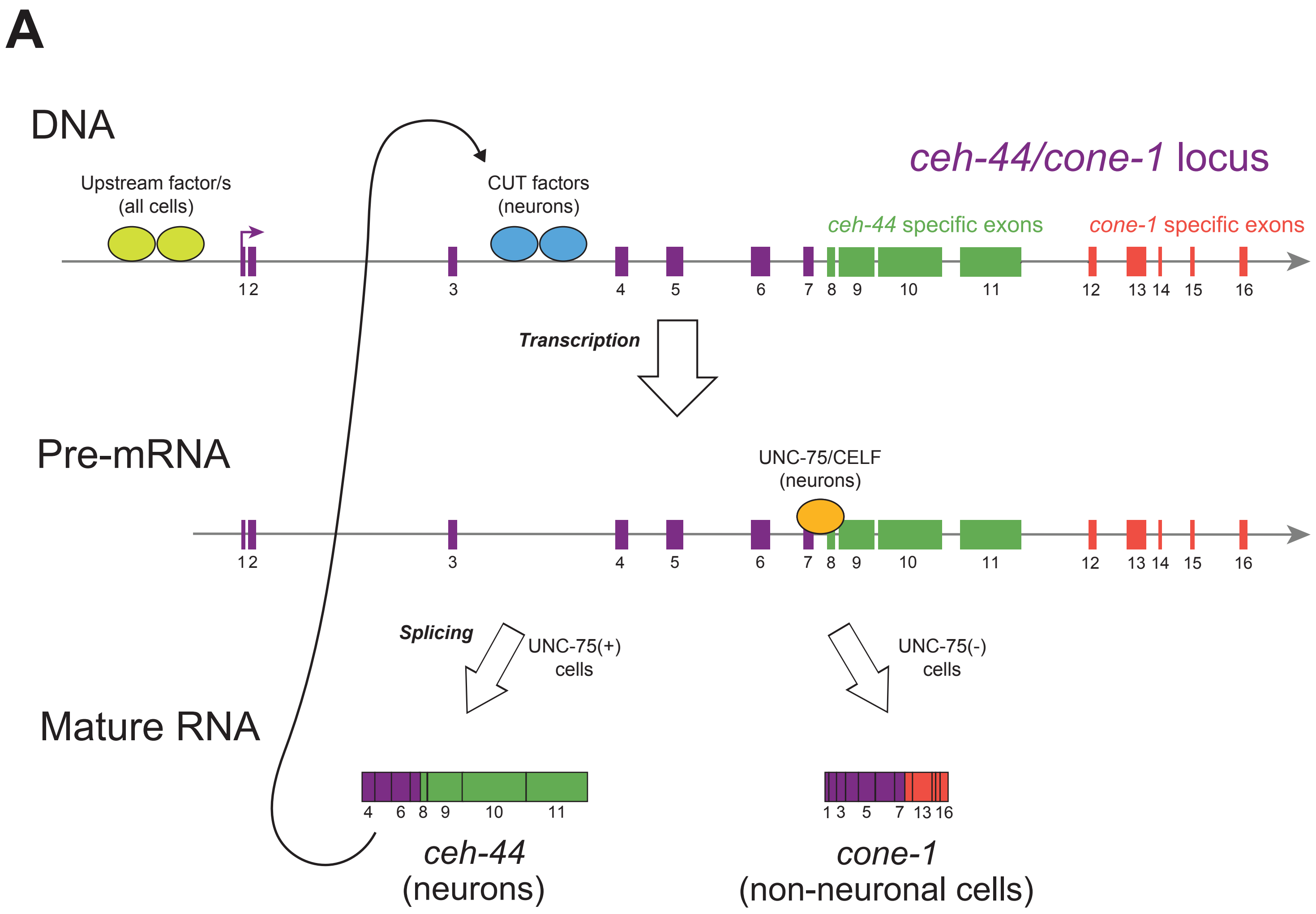
A
DNA
ceh-44/cone-1 locus
Upstream factor/s
(all cells)
CUT factors
(neurons)
ceh-44 specific exons
cone-1 specific exons
1 2
Transcription
Pre-mRNA
UNC-75/CELF
(neurons)
1 2
Splicing
UNC-75(+)
cells
UNC-75(-)
cells
Mature RNA
4
6
8
9
10
11
1 3
5
7
13 16
ceh-44
(neurons)
cone-1
(non-neuronal cells)
